# The effect of carnitine on preservation of renal cold ischemia in rats

**DOI:** 10.1101/494963

**Authors:** Arif Aslaner, Ömer Günal, Hamdi Taner Turgut, Ümran Yıldırım

## Abstract

**Background:** Although, University of Wisconsin (UW) solution is referred to as the gold standard for the preservation from cold ischemia, studies that aiming to protract the cold ischemia time still goes on. This study was planned to evaluate the effectiveness of carnitine that is added to UW solution on renal cold ischemia time.

**Methods:** Thirty two male Wistar Albino rats were divided into four groups of 8 each; Control, UW, carnitine with Ringer Lactate (RL) and UW groups. Retrieved renal grafts were preserved in solutions at +4°C after perfusion. At the same time tissue samples of 72th hour was assessed to measure Malondialdehyde (MDA) levels. Preservation solution samples were assessed at 0th, 24th, 48th, and 72th hours to measure Lactate Dehydrogenase (LDH) activity. Tissue injury was histologically scored by evaluating the hematoxilen-eosin stained paraffin sections.

**Results:** Administration of carnitine to preservation solutions decreased the mean tissue MDA levels in groups 2 and 4. Carnitine administration slowed the increase of LDH levels in preservation solutions as well as the histopathological total tissue injury scores.

**Discussion:** We conclude that the addition of carnitine to UW solution increased the preventive effect of preservative solution on cold ischemia injury of kidney.

## 1. Introduction

Cold ischemic organ preservation is the most commonly used procedure for almost all solid organs. Despite University of Wisconsin (UW) solution is the most commonly used and gold standard solution for the cold organ preservation of kidney, further investigations still continues. Since the beginning of the 1990s, UW solution has been used in heart, lung, liver, pancreas, kidney, and intestinal transplants with great success. Using UW solution for preservation, the ischemia tolerance limit has been extended to 18 h for liver and 36 h for kidney1. In the prevention of injury after ischemia many antioxidants and free radical scavengers such as carnitine, a water-soluble quaternary ammonium structure and naturally present in the cells of animal origin, have protective effect on renal ischemia injury2,3. Carnitine also prevents early graft dysfunction4 and is partially preventing renal damage developed as a result of ischemia reperfusion5. This present study was performed to investigate the any actual benefit of the carnitine alone or in conjuction with other cold preservation solutions.

## 2. Material and Methods

### Animals

This study was conducted at the Physiology Laboratory with thirty two male Wistar albino rats weighing 225–250 gr and aging 12-18 m. The principles of laboratory animal care were strictly followed. Rats were taken to laboratory unit, placed in controlled conditions of temperature (23±2°C) and 12/12-hour light-dark cycles. And they acclimatized for 2 weeks adhering to the ethics of research on animals and fed with standard rat chow and water ad libitum. Food intake was withheld overnight before surgery. Rats were randomly and equally divided into four groups. Standard surgical model procedure was performed to all groups.

### Preservation Solutions and study groups

Specimens were preserved in 40ml Ringer lactate (RL) solution in group one (n=8); in 40ml RL solution containing carnitine in group two (n=8); in 40ml UW solution in group three (n=8) and in 40ml UW solution containing carnitine in group four (n=8). The 22 mg/ml carnitine was added to Groups 2 and 4. All groups were also perfused before renal harvesting with preservation solutions in which their specimens have been preserved later.

### Anesthesia and Experimental Design

Following a 24 hours of starvation, the rats were anesthetized with intraperitoneal ketamine (Ketalar flacon-10 mg/kg, Pfizer Warner Lambert) and chlorpromazine (Largactil 25mg ampule-0,1mg/kg, Eczacibasi Rhone-Poulenc). They were weightened and operated randomly. The abdominal cavity was opened via midline incision. The aorta, inferior vena cava (IVC), left and right renal vessels were atraumatically isolated. The infrarenal abdominal aorta was cannulated with 23-gauge polyethylene catheter tube and the tip of the catheter set forward above renal artery level. Then the IVC was catheterized to enable the exanguination of blood and perfusion solution. Two milliliters of heparinized physiological saline were infused through the abdominal aorta and two milliliters of blood sample were taken for biochemical analysis.

Kidneys were perfused with the preservation solutions in which they would be preserved in, at +4oC through the aorta continuously at 70cmH2O pressure until the both kidneys paled and the output from the IVC cannula was as clear as the perfusate. Finally, bilateral nephrectomy was performed. All the graft tissues were immersed and preserved in preservation solution immediately after harvesting. The grafts tissues were weightened and kept in sterile plastic containers containing 40ml preservation solution at +4oC in a refrigerator. One milliliter of a mixed sample of preservation fluid was taken from the container for enzyme measurements at 0, 24, 48, and 72 hours.

### Biochemical Analysis

The LDH activities in the serum and preservation solutions at 0, 24, 48, and 72 hours were determined by standard clinical chemistry methods using an Olympus AU 640 autoanalyzer (Olympus, Tokyo, Japan).

### Measurement of malondialdehyde levels

To determine the MDA levels, kidney tissue samples were homogenized in ice cold 150mm KCl. The MDA levels (nmol MDA/g tissue) were assayed for the products of lipid peroxidation. Tissue MDA levels were measured at 72th hour using the method described by Uchiyama et al6.

### Histopathological Evaluation

Renal injury was analyzed and reviewed by a pathologist who is blind to the groups and preservation solutions. Samples were fixed and embedded in hematoxylin–eosine (H&E) stained paraffine sections for examination under light microscopy (Olympus BX 50). Samples were also examined for cellular ultrastructure under an electron microscope (JEOL, Tokyo, Japan).

Light microscopic sections were examined for vacuolization, denuded basement membrane, necrosis, tubular dilatation, and cell detachment. Electron-microscopic sections were examined for mitochondria and tubular cell brush border integrity and interstitial and cell edema. Nine basic morphological patterns (apical cytoplasm vacuolization, tubular necrosis, denuded basement membrane, tubular dilatation, cell detachment, intracellular edema, interstitial edema, brush border integrity, mitochondria integrity) were graded on a five-point scale as following: 1, no abnormality; 2, mild lesions affecting 10% of kidney samples; 3, lesions affecting 25% of kidney samples; 4, lesions affecting 50% of kidney samples; and 5, lesions affecting more than 75% of kidney samples7.

### Statistical Analysis

ANOVA was performed by using SPSS 15.0 (Statistical Package for Social Sciences; SPSS Inc, Chicago, Illinois, USA) for Windows and Microsoft Office Excel 2010 version for evaluating the data. In comparison the data Kruskal Wallis, Friedman, Mann Whitney U, Wilcoxon, Chi square and Fisher’s Exact Tests analysis were all used. For descriptive anlysis of the data, numerical variables and minimum maximum values were expressed as means ± standard deviation (SD). Values of P < 0.05 were considered as statistically significant.

## 3. Results

In the present study, the LDH levels in the serum and preservation solutions of all groups at 0th, 24th, 48th, and 72th hours and MDA levels at 72th hour were measured. In all groups, before perfusion two milliliters of blood were taken from abdominal aorta to evaluate the renal function. Despite the LDH levels of preservation solutions at 0th and 24th hours of group 1 and 2 were lower than the group 3 and 4. At 48th and 72th hours a further increase was observed in group 1 and 2 than group 3 and 4 (Fig. 1). The MDA levels of group 1 was higher than groups 2,3 and 4. According to tissue MDA levels, there is a significant difference between all groups except group 2 and group 3. (Table 1)

**Fig 1:**
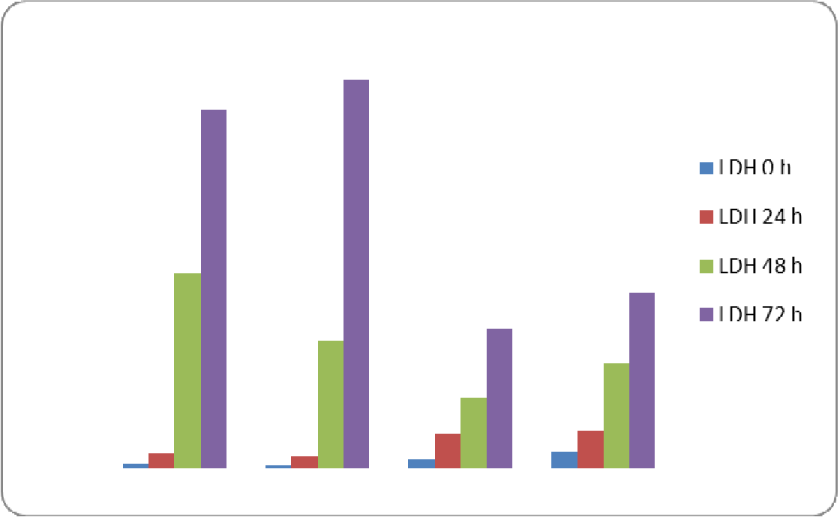
Distribution of mean LDH levels of preservation solutions at 0, 24, 48, 72 hours.

**Table 1:**
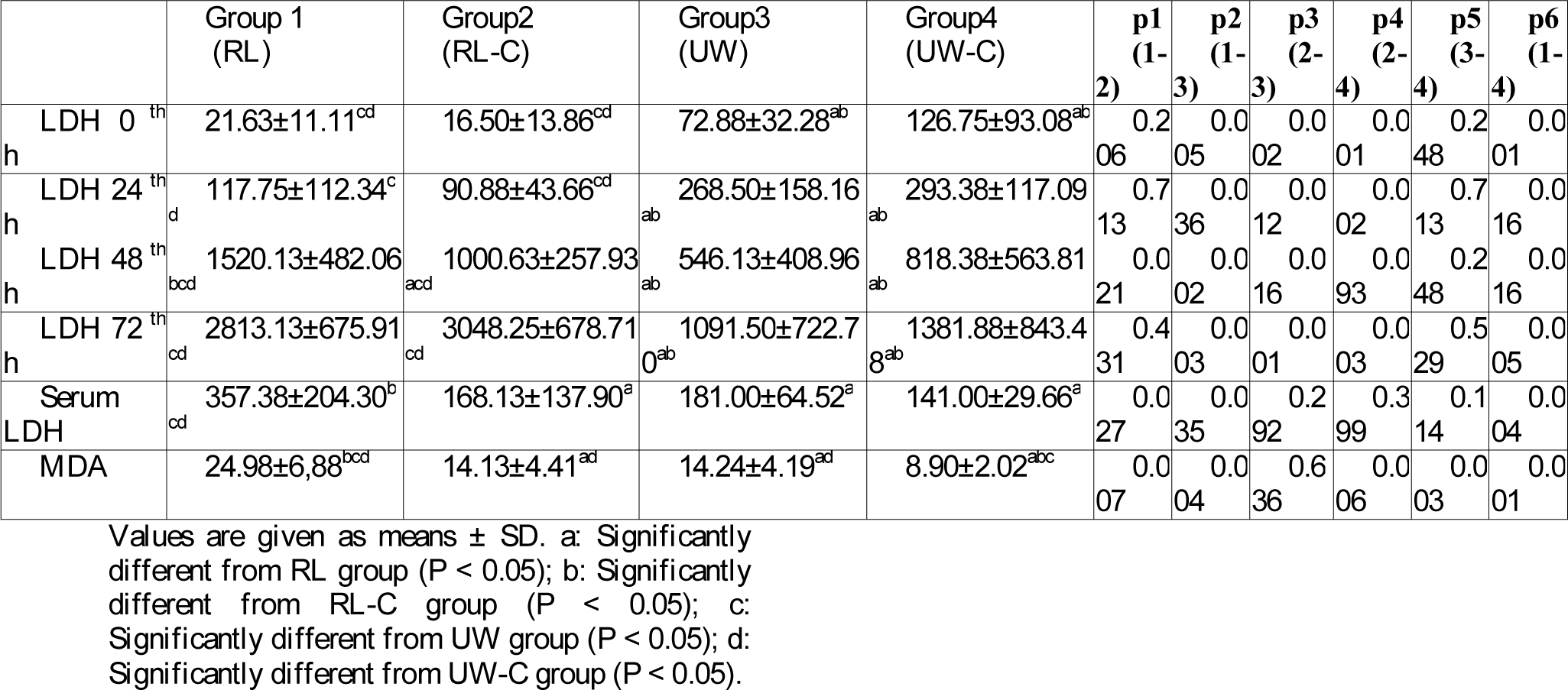
Tissue MDA (nmol/gr tissue) and LDH (IU/L) levels of serum and preservation solutions of groups.

In the histopathological evaluation of the samples; apical cytoplasm vacuolization, tubular necrosis, denuded basement membrane, tubular dilatation, cell detachment, intracellular edema, interstitial edema, brush border integrity and mitochondria integrity were all significantly decreased in groups 3 and 4 than groups 1 and 2. (Table 2)

**Table 2:** The histopathological injury scores of the groups.

|  | Group 1<br>(RL) | Group2<br>(RL-C) | Group3<br>(UW) | Group4<br>(UW-C) | p1<br>(1-2) | p2<br>(1-3) | p3<br>(2-3) | p4<br>(2-4) | p5<br>(3-4) | p6<br>(1-4) |
| --- | --- | --- | --- | --- | --- | --- | --- | --- | --- | --- |
| **Apical cytoplasm vacuolization** | 2.25±0.46<sup>cd</sup> | 2.5±0.53<sup>cd</sup> | 1<sup>ab</sup> | 1<sup>ab</sup> | 0.608 | < 0.001 | < 0.001 | < 0.001 | 1 | < 0.001 |
| **Tubular necrosis** | 4.5±0.53<sup>bcd</sup> | 2.5±<sup>acd</sup> | 1<sup>ab</sup> | 1<sup>ab</sup> | 0.003 | < 0.001 | 0.002 | 0.418 | 0.064 | 0.012 |
| **Denuded basement membrane** | 3<sup>bcd</sup> | 1.88±0.35<sup>acd</sup> | 1<sup>ab</sup> | 1<sup>ab</sup> | < 0.001 | < 0.001 | 0.001 | 0.001 | 1 | < 0.001 |
| **Tubular dilatation** | 3.63±0.52<sup>bcd</sup> | 2.38±0.74<sup>acd</sup> | 1<sup>ab</sup> | 1.13±0.35<sup>ab</sup> | 0.027 | < 0.001 | 0.002 | 0.009 | 1.000 | 0.001 |
| **Cell detachment** | 4±0.53<sup>bcd</sup> | 2.38±0.52<sup>acd</sup> | 1.13±0.35<sup>ab</sup> | 1<sup>ab</sup> | 0.005 | 0.003 | 0.002 | < 0.001 | 1.000 | 0.001 |
| **Intracellular edema** | 3.88±0.35<sup>bcd</sup> | 1.75±0.71<sup>acd</sup> | 1<sup>ab</sup> | 1<sup>ab</sup> | 0.003 | < 0.001 | 0.026 | 0.026 | 1 | < 0.001 |
| **Interstitial edema** | 3<sup>bcd</sup> | 3<sup>acd</sup> | 1<sup>ab</sup> | 1<sup>ab</sup> | 0.315 | < 0.001 | < 0.001 | 0.018 | 0.200 | 0.007 |
| **Brush border integrity** | 4.88±0.35<sup>cd</sup> | 3±0.53<sup>cd</sup> | 1.25±0.46<sup>ab</sup> | 1.25±0.46<sup>ab</sup> | 0.003 | 0.001 | 0.004 | 0.004 | 1.000 | 0.001 |
| **Mitochondria integrity** | 2<sup>bcd</sup> | 2<sup>acd</sup> | 1<sup>ab</sup> | 1<sup>ab</sup> | 0.006 | 0.001 | < 0.001 | 0.005 | 0.077 | 0.003 |

Values are given as means ± SD. a: Significantly different from RL group ( $P < 0.05$ ); b: Significantly different from RL-C group ( $P < 0.05$ ); c: Significantly different from UW group ( $P < 0.05$ ); d: Significantly different from UW-C group ( $P < 0.05$ ).

## 4. Discussion

Contemporary approach to organ transplantation necessitates three main goals; utilize appropriate organs for transplantation, improve and pay attention for the initial functions after organ transplantation and the convenient and trusted organ protection7. Initial organ function after transplantation is mainly related to the surgical capabilities, the preservation method and the cost-effectiveness of preservation method8.

Clinically, the renal preservation with continuous hypothermic perfusion9 and simple cold preservation10 has been developed towards the end of 1960.

There are three main organ preservation; hypothermia, to prevent cell swelling and the biochemical effects8. It has been shown that simple cold preservation of the kidneys were effective 72 hours experimentally, and 24 to 50 hours clinically2. Hypothermia reduces the metabolic requirements of the tissues, and thus allows reliable protection than normothermic conditions11.

Hypothermia inactivate sodium potassium pump and if this status is not corrected this swelling leads to cellular injury11.

In many clinical conditions, kidneys and the other organs in the body may expose to ischemia. Cells interfere with renal tubular damage in renal ischemia. Shock and kidney transplants are the most common clinical conditions that interfere with renal ischemia. Treatment option for end-stage renal failure is renal transplantation. In the prevention of injury after ischemia, many antioxidants and free radical scavengers (α-tachopherol, allopurinol, glutathione, superoxide dismutase, desferroxamine, etc.) have protective effects12.

Carnitine is another agent used for many series of clinical and experimental studies that have shown the effects of anti-ischemic effects on myocardium, and skeletal muscle. Carnitine is a water-soluble quaternary ammonium structure, a compound oftenly naturally present in the cells of animal origin. Carnitine in the diet is quickly absorbed into the intestinal lumen of a place of passive and active mucosal transport. At normal ranges 98% of the total carnitine pool was at heart and skeletal muscle, 1.6% at the liver and kidney, and the remaining 0.6% is located in the extracellular fluid13.

Carnitine is an antioxidant that prevents the accumulation of lipid peroxidation end-products. Carnitine is an essential amino acid that syntesized from two non-essential amino acid lysine and methionine in liver and kidney. And they transported from main production sites liver and kidneys to skeletal and cardiac muscle. At the same time metabolites of carnitine acyl-CoA is important for beta-oxidation of free fatty acids by transport from the mitochondrial inner membrane to matrix14,15.

Some researchers also found that carnitine is effective in renal ischemia injury2,3, prevent early graft dysfunction4 and is partially preventing renal damage developed as a result of ischemia reperfusion5. Carnitine is also combinated with 5-aminoimidazole-4-carboxyamide ribonucleoside that enhancing energy metabolism and provides a novel modality to treat renal ischemia/reperfusion injury16. Some researchers were demonstrated that carnitine provide in vitro and in vivo membrane fluidity and also prevent oxygen free radicals13.

In a study that metabolically active substrate carnitine was added to preservation solutions, a significant increase in metabolism by increasing the ATP levels and good ultrastructural integrity of mitochondria and endoplasmic reticulum shown to provide significant protection. They conclude that carnitine significantly increase the graft viability after preservation17.

In our study we observed that LDH levels of preservation solutions at 0h and 24h of groups 1 and 2 were lower than the groups 3 and 4. And also at 48th and 72th hours a further increase on LDH levels were observed in groups 1 and 2 than groups 3 and 4.

According to these results in order to preserve the organs from cold ischemic injury for 24 hours the RL and carnitine containing RL solutions can be perceived as adequate. If long term preservation needed such as 48 to 72 hours or more, UW and carnitine containing UW solution would be better.

In our present study we found significant differences on tissue malondialdehyde levels between groups 1 with 2 and groups 3 with 4. (p=0,001; p=0,005 respectively). According to this result we demonstrated that adding carnitine to preservation solutions resulted in a decrease in tissue MDA levels. There are many studies that offer similar results to our findings3,4,12,13,15,18.

In histopathological examination nine basic morphological patterns as follows; apical cytoplasm vacuolization, tubular necrosis, denuded basement membrane, tubular dilatation, cell detachment, intracellular edema, interstitial edema, brush border integrity, mitochondria integrity were graded on a five-point scale were evaluated by an experienced pathologist. Although there is not any difference between group 3 and group4, there is a significant reduction on injury in carnitine containing group 2 according to group 1.

Nowadays hypothermic ischemic preservation is the commonly and effectively used method for a period of ischemic organ protection. The most important parts of this technique are hypothermia and chemical solutions. Despite all these, to provide longer period of organ protection time further studies are needed.

## 5. Conclusions

In this present study we aimed to investigate the effect of carnitine on ischemic injury of kidney by adding it to preservation solutions. We found that adding carnitine to preservation solutions prolongs the storage time and may contibute to preservative function.

As a result of these findings, we concluded that if the grafts were planned to protect for longterm storage such as 48 and 72 hours carnitine adjunction to preservative solutions such as RL and UW may be able to prolong the preservation time effectively.

